# Novel genotyping algorithms for rare variants significantly improve the accuracy of Applied Biosystems ™ Axiom ™ array genotyping calls

**DOI:** 10.1101/2021.09.13.459984

**Authors:** O Mizrahi Man, MH Woehrmann, TA Webster, J Gollub, A Bivol, SM Keeble, KH Aull, A Mittal, AH Roter, BA Wong, JP Schmidt

## Abstract

**Objective:** To significantly improve the positive predictive value (PPV) and sensitivity of Applied Biosystems™ Axiom™ array variant calling, by means of novel improvement to genotyping algorithms and careful quality control of array probesets. The improvement makes array genotyping more suitable for very rare variants.

**Design:** Retrospective evaluation of UK Biobank array data re-genotyped with improved algorithms for rare variants.

**Participants:** 488,359 people recruited to the UK Biobank with Axiom array genotyping data including 200,630 with exome sequencing data.

**Main Outcome Measures:** A comparison of genotyping calls from array data to genotyping calls on a subset of variants with exome sequencing data.

**Results:** Axiom genotyping [18] performed well, based on comparison to sequencing data, for over 100,000 common variants directly genotyped on the Axiom UK Biobank array and also exome sequenced by the UK Biobank Exome Sequencing Consortium. However, in a comparison to the initial exome sequencing results of the first 50K individuals, Weedon et al. [1] observed that when grouping these variants by the minor allele frequency (MAF) observed in UK Biobank, the concordance with sequencing and resulting positive predictive value (PPV) decreased with the number of heterozygous (Het) array calls per variant. An improved genotyping algorithm, Rare Heterozygous Adjustment (RHA) [16], released mid-2020 for genotyping on Axiom arrays, significantly improves PPV in all MAF ranges for the 50K data as well as when compared to the exome sequencing of 200K individuals, released after Weedon et al. [1] performed their comparison. The RHA algorithm improved PPVs in the 200K data in the lowest three frequency groups [0, 0.001%), [0.001%, 0.005%) and [0.005%, 0.01%) to 83%, 82% and 88%; respectively. PPV was above 95% for higher MAF ranges without algorithm improvement. PPVs are somewhat higher in the 200K dataset, due to a different “truth set” from exome sequencing and because monomorphic exome loci are not included in the joint genotyping calls for the 200K data set, as explained in the methods section.

Sensitivity was higher in the 200K data set than in the original 50K data as well, especially for low MAF ranges. This increase is in part due to the larger data set over which sensitivity could be computed and in part due to the different WES algorithms used for the 200K data [7]. Filtering of a relatively small number of non-performing probesets (determined without reference to the exome sequencing data) significantly improved sensitivities for all MAF ranges, resulting in 70%, 88% and 94% respectively in the three lowest MAF ranges and greater than 98% and 99.9% for the two higher MAF ranges ([0.01%, 1%), [1%, 50%]).

**Conclusions:** Improved algorithms for genotyping along with enhanced quality control of array probesets, significantly improve the positive predictive value and the sensitivity of array data, making it suitable for the detection of very rare variants. The probeset filtering methods developed have resulted in better probe designs for arrays and the new genotyping algorithm is part of the standard algorithm for all Axiom arrays since early 2020.

## Introduction

UK Biobank [3] is a highly valuable health resource that follows the health and well-being of 500,000 participants, all genotyped on Thermo Fisher Scientific’s Applied Biosystems™ Axiom™ microarrays. Whole exome sequencing (WES) data for a subset of participants was also made available [6,7]. The initial WES release contained data for about 50,000 participants [6]. In a manuscript recently published, Weedon et al. [1] compared the genotyping calls from the microarray to those in the WES data from this initial release, as a function of minor allele frequency (MAF) computed from the array calls (cMAF). They observed that heterozygous concordance is very high in cMAF ranges above 0.01% but drops off for a cMAF range between 0.001% and 0.01% and drops further for cMAF ranges below 0.001%. This lowest MAF group corresponds to variants with fewer than 9 individuals out of ~500,000 with non-homozygous array genotypes.

Very rare variants can be more difficult to genotype on microarray platforms because they use clustering algorithms to identify the genotype [8,9,13]. While for common variants the location and shape of the heterozygous (Het) cluster provides powerful evidence for the accuracy of Het calls, rare variants often have no other samples in the Het cluster, making the call less robust.

In-depth analysis of the distribution of replicate probe signals on Axiom microarrays revealed that this distribution can be used to detect false Het calls with remarkable accuracy. We studied the nature of false positive array calls using 1000 Genomes Project [2] reference samples on a variety of Axiom arrays and developed an algorithm specifically targeted to improve rare Het calls. Here we present the Rare Het Adjusted (RHA) Algorithm, a novel genotyping algorithm, that eliminates the majority of false positive heterozygous calls, while leaving true positive heterozygous calls virtually untouched. RHA is a relatively small addition to the Axiom™ GT1 algorithm [8], tailored to very rare variants, that significantly improves PPV for variants with MAF below 0.01%. Higher MAF ranges have excellent PPV (above 95%) even without RHA (Table 1). In 2021, WES data for an additional 150,000 participants was released; we compared our improved algorithms to both the initial exome data [6] and the combined exome data for 200,000 individuals [7]. Results vary depending on the set that is used as truth (50K exome sequencing versus 200K exome sequencing data) and based on the quality control applied to WES data. While PPV always improves when stricter quality control is applied to the sequencing data, the conclusions in Weedon et al. (prior to algorithm improvements) hold up directionally in the 200K exome data, as do the significant improvements following RHA. The results presented here did not apply stricter quality control to the WES data. Sensitivity is higher in the 200K dataset in all minor allele frequency ranges, likely due to the larger data set over which sensitivity can be computed and due to different WES genotyping algorithms applied to the 200K data vs the 50K data. Filtering of a relatively small number (2,215) of non-responsive probesets (identified without using the exome sequencing data), significantly improved sensitivities for all MAF ranges. The probeset filtering methods developed have resulted in better array probe design practices, and the new RHA algorithm is part of the standard algorithm for all Axiom arrays since early 2020.

**Table 1.**
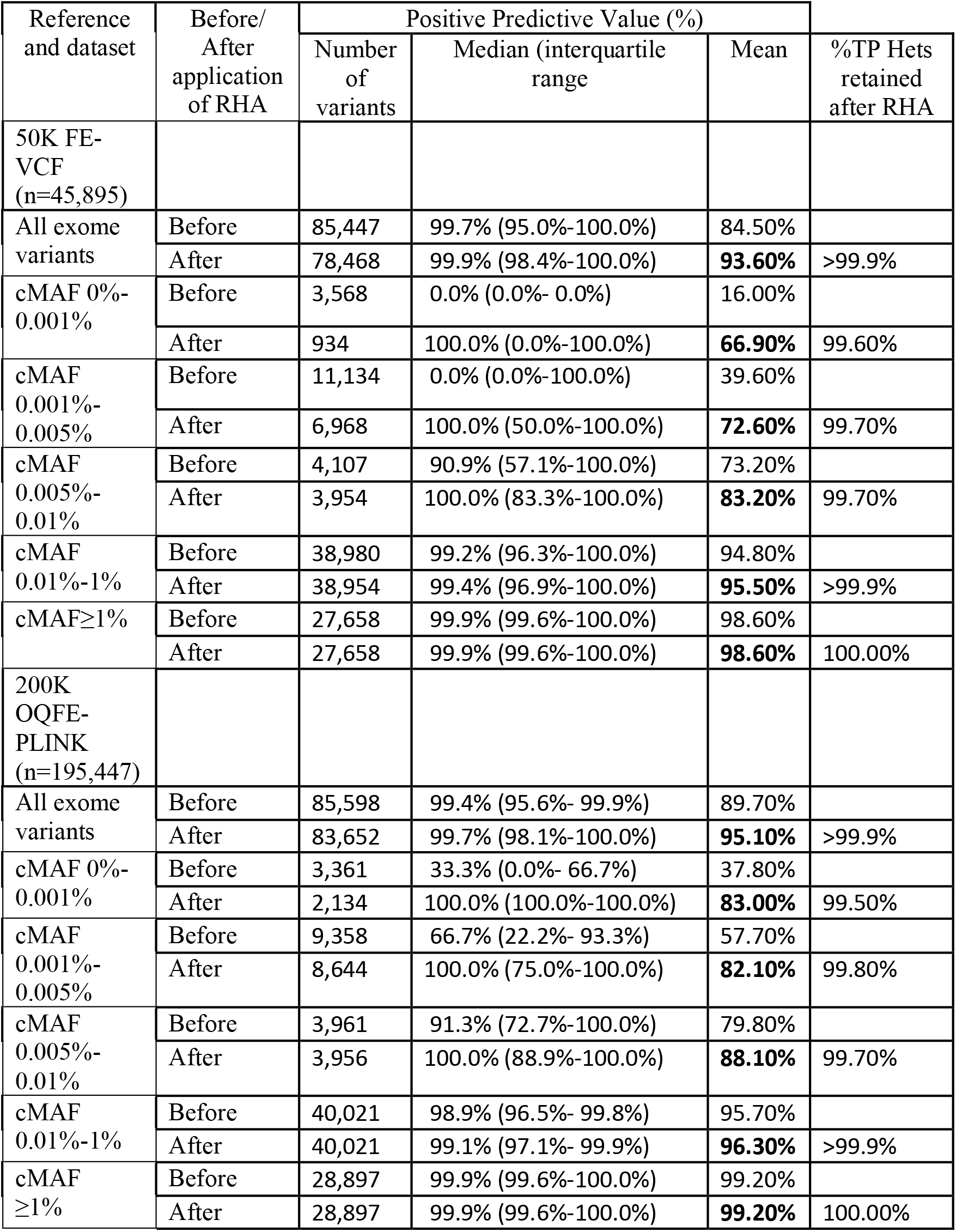
Positive predictive value of UK BioBank Axiom™ array versus whole exome sequencing before and after application of RHA. Results are split by the WES dataset used as reference - 50K FE-VCF and 200K OQFE-PLINK, and by the minor allele frequency calculated from the array before application of RHA (cMAF). We include the following summary statistics, calculated across all relevant single nucleotide variants: median and inter-quartile range of PPV (the distribution of PPV is not normal) as well as the mean PPV, for the purposes of comparison with Weedon et al. Also included are the number of variants the summary statistics have been calculated over (only variants with at least one TP or FP genotype can yield a PPV) and the percentage of TP Hets that are retained after applying RHA measured as (TP Hets post RHA)/(TP Hets pre RHA) (only Het genotypes are affected by the algorithm). Insertions and deletions have been excluded from calculations. The lower PPV (pre RHA) in the 50K FE-VCF as compared to the 200K OQFE data set is largely due to the many loci included in the 50K FE-VCF that are monomorphic in the Exome data. Since RHA is very efficient at eliminating FP Hets, the difference diminishes significantly post RHA

## Methods

### The Rare Het Adjusted (RHA) algorithm

The Rare Het Adjusted algorithm (RHA) is executed after the original genotyping algorithm (AxiomGT1), to determine which predictions of rare heterozygous (Het) genotypes are likely to be false. The genotypes of false Het predictions, as determined by RHA, are then set to ‘No Call’. The main insight of the algorithm, after examining many hundreds of different array results, is that most false Het predictions occur due to a small number of higher or lower than expected intensities on the microarray surface. These unexpected intensities cause a homozygous sample’s summarized probeset intensity to be positioned away from the major homozygous genotype cluster in “signal contrast” vs. “signal size” (clustering) space and into a location that could be consistent with a heterozygous genotype cluster, thereby causing the AxiomGT1 genotyping algorithm to falsely predict a Het.

The AxiomGT1 algorithm used to genotype the UK Biobank already incorporates detection and elimination of unexpected intensities into its genotype calling but does not use higher stringency for low MAF variants. However, genotyping variants for which Hets are rare in the population is especially challenging because false Het predictions are more likely to occur in cases where the number of true Hets is too low (or zero in the most extreme case) to correctly anchor the Het cluster [1,14]. RHA improves rare Het prediction by incorporating additional information in the test for unexpected intensities. These include: (1) the intensity differences of replicated probe sequences on the array for the putative Het sample and (2) the position of the putative Het signal relative to the signal distribution of homozygous samples.

RHA is summarized in Fig. 1. The algorithm is applied to probesets with two replicate probes for bi-allelic variants and a small number of Hets. In the UK Biobank release samples were genotyped in batches of about 5,000 samples and RHA is applied only if the number of Het predictions in the particular batch is between 1 and max_nAB. Max_nAB is a parameter of the algorithm; in our analysis we used a value of 4.

**Fig. 1.**
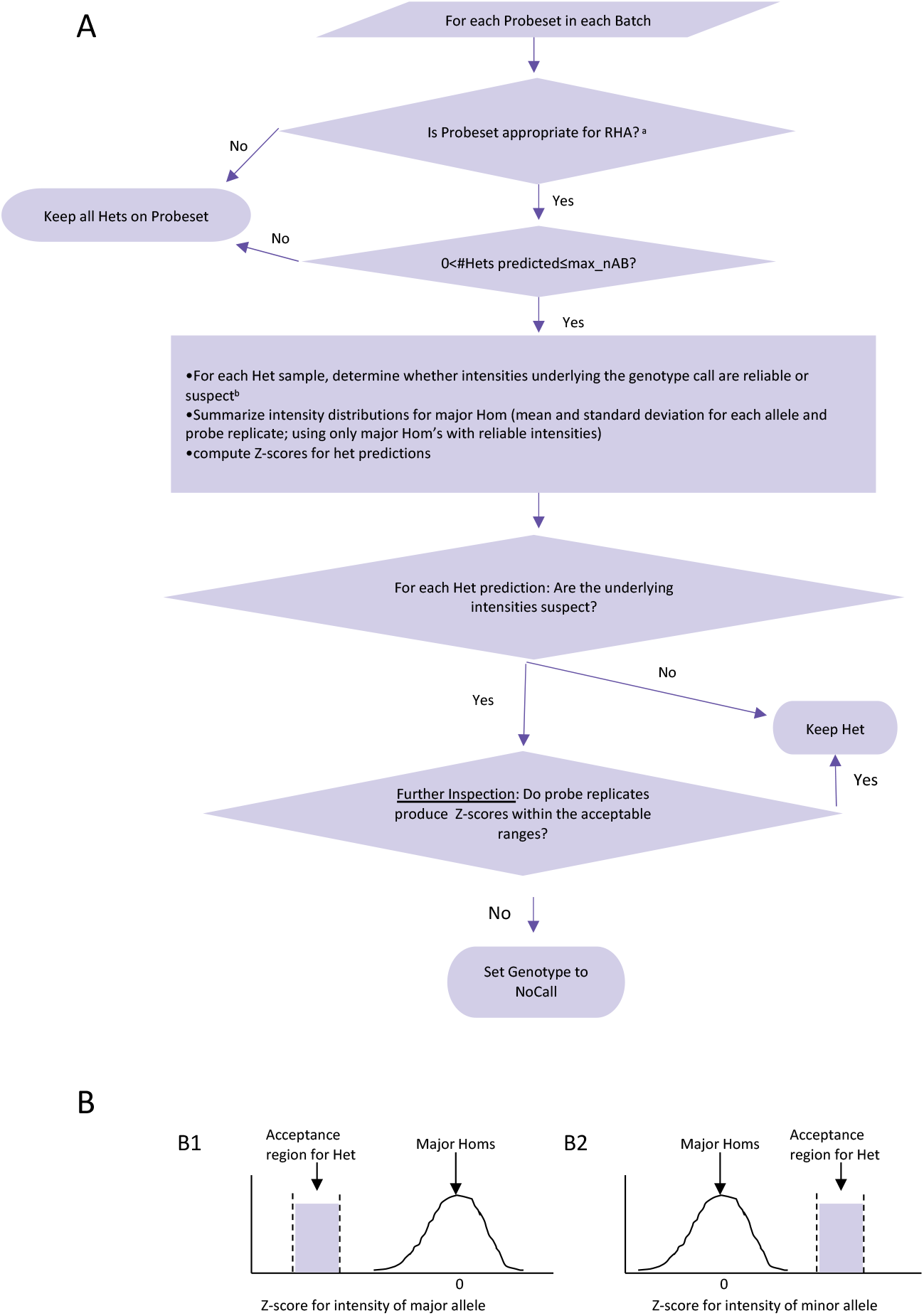
RHA scheme for distinguishing true Hets from false ones. This is a post-processing scheme that accepts as input the genotypes called by AxiomGT1, the normalized intensities and the annotation of which locations on the array were eliminated by AxiomGT1 before calling genotypes. A. overall flow of the algorithm; B. Illustration of acceptable range for Het intensity Z-scores relative to the distribution of intensities from major Hom samples. Z-scores for the intensities of predicted Hets are calculated, based on summary statistics (mean and standard deviation) of the distribution of corresponding intensities for major Homs on the particular probeset, using the following formula: 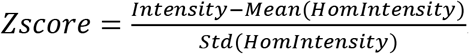. We use only major Hom genotypes with reliable underlying intensities to calculate these summary statistics. In addition, we compute Z-scores only when the major allele can be determined with confidence (at least *90%* of samples are major Hom) and number of high-quality major Hom genotypes is sufficiently high (at least *30*). Finally, if we were unable to obtain summary statistics for major Homs, all Hets with underlying aberrant intensities are set to ‘No Call’. B1. Acceptance region for the Z-score of the major allele intensity. Since major Hom’s have two copies of the major allele, whereas a Het has only one copy of this allele the Z-score is expected to be negative. However, intensity values and their associated Z-scores that are too low may be indicative of a probe that isn’t working and are thus not accepted. B2. Acceptance region for the Z-score of the minor allele intensity. Major Hom’s have zero copies of the minor allele, and their intensity on this allele is thus the background noise, whereas a Het has one copy of this allele. Thus, we expect the Z-score for the minor allele intensity to be positive. On the other hand, a Z-score that is too high could indicate that the intensity measured on the probe is aberrantly high. ^a^ A probeset is appropriate for RHA if it is bi-allelic, two replicates on the array and a small number of Hets. ^b^ The intensities for a specific probeset and sample are deemed reliable if AxiomGT1 used both replicates in genotyping and SCI is low for both alleles; otherwise, the intensities are deemed suspect and so is the resulting genotype call.

All Het calls for rare variants are tested for patterns of “unexpected intensities” which are defined as an inconsistency in intensities between the two replicate probe sequences for the Het. If there are no unexpected intensities, the prediction is deemed to be true, and the Het genotype call is retained. Otherwise, the prediction undergoes further inspection to determine whether the individual probe intensities are “Het-like”. If the prediction passes this second inspection, the Het genotype call is retained, otherwise the genotype of the probeset on the sample is set to “No Call” (NC).

### Suspect Het predictions with unexpected intensities

The AxiomGT1 genotyping algorithm clusters samples in “signal contrast” vs. “signal size” (clustering) space to obtain genotypes, after summarizing the intensities for the various replicates of the same probe [8]. Different replicates of probe sequences for the same probeset may have a slightly different intensity distribution as a result of their locations on the array. Minor non-niformities on the array surface can cause a probe’s intensity to be lower or higher than it should be.

With RHA (the improved algorithm), a Het prediction is marked as requiring further inspection, if it is a rare Het (Het cluster size <= max_nAB) and one of two criteria for an unexpected intensity is met:

a. **One of the probeset’s probe replicates was eliminated by the Axiom GT1 algorithm because of an unexpected intensity**. For such probes, the genotyping algorithm uses just one of the replicates to genotype the sample. However, on occasion the single remaining intensity may cause a shift in the position of a homozygous sample in clustering space that is sufficiently large and in an appropriate direction for the genotyping algorithm to call a Het.
b. **The difference between intensities measured on different probe replicates is higher than expected.** To predict and classify probe intensities as unexpected, we devised the SCI (Sequence Contrast Index) metric. SCI is calculated separately for the A and B alleles, using I1 and I2, the normalized intensities for the two probe replicates and can easily be generalized for more replicates:

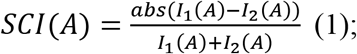

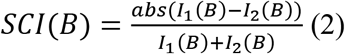

SCI equals 0 for identical intensities and approaches 1 as the difference between intensities grows. Because both replicates of a probe are unlikely to experience the same type of unexpected intensity at the same time, a low SCI indicates that intensities measured on both probe replicates are reliable. If SCI is high for at least one of the alleles, the Het prediction is marked for further inspection. A cutoff value for SCI was empirically derived and has been shown to be applicable for different arrays and batch sizes.

### Further inspection of suspect Het predictions

Not all Het predictions with unexpected underlying intensities are false. To minimize loss of true positive Hets, we further examined suspect Het predictions. Ideally, we would compare the intensities driving the Het prediction to a distribution of expected intensities for a Het genotype, as is done for higher MAF variants. However, this requires many examples of true Hets for the given variant, which are generally not available for low MAF frequency variants. Instead, we compare the predicted Het’s intensities to the distribution of intensities for the major Hom. A major Hom genotype is comprised of two copies of the major allele and zero copies of the minor allele, whereas a Het genotype is comprised of one copy of each allele. These differences should be reflected in the intensity measurements of each of the two alleles and can therefore be used to differentiate the two genotypes. The advantage of this approach is that for low minor allele frequencies the vast majority of genotypes are major Homs, giving us plenty of examples to characterize the major Hom intensity distribution for a given replicate. While the intensity distribution differs for different probesets, and sometimes for different replicates, we have found that the range of positions relative to the mean and variance of this distribution that is appropriate for a Het is universal for all probesets. We therefore translate the intensities underlying the Het prediction into Z-scores relative to the appropriate major Hom intensity distribution and check whether these Z-scores fall in the “acceptable” range (different ranges are appropriate depending on whether the intensity is measured for the major or minor allele). In this way we check for each probe-replicate whether the intensities measured on it are uniformly Het-like (Fig. 1B).

### Derivation of algorithm parameters

To select appropriate values for RHA parameters, we trained on six independent datasets, each comprising one to three array plates and containing 1000 Genomes Project [2] reference samples, using sequencing data from the 1000 Genomes Project as truth. Training was limited to putative singleton Hets (i.e., a cluster size of one). Balancing specificity and sensitivity, we selected, using a simple grid search of parameter space, a cutoff for SCI, as well as minimum and maximum Z-scores (data not shown). It is worth noting that as the number of Hets per cluster increases, fewer Het genotypes have unexpected signals and are set to NC by RHA (Fig. 2).

**Fig. 2.**
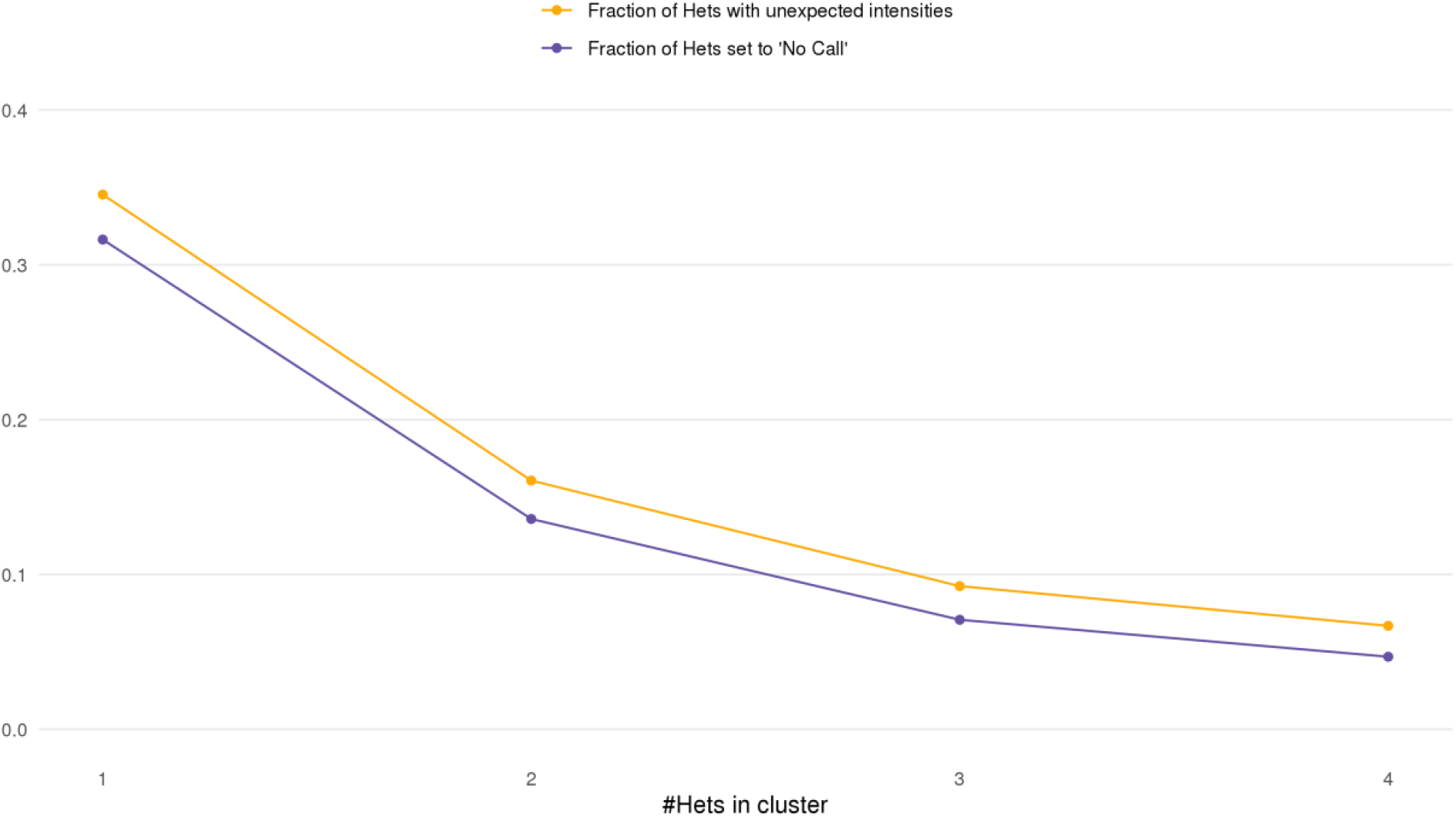
The fraction of predicted Hets that are set to ‘No Call’ by RHA decreases as the size of the Het cluster increases. For cluster sizes ranging from 1 to 4, we show the fraction of predicted Hets with underlying unexpected intensities and the fraction of Hets that are set to ‘No Call’. Both quantities decrease with increasing size of Het cluster.

### Axiom genotyping data

The analyses in this paper center around the genotyping data for samples genotyped on the Applied Biosystems™ UK Biobank Axiom™ array for which WES data is available. The results for genotyping data generated on the UK BilEVE array for which WES data is available are presented in the supplementary material and comprises only 3% of the samples with WES. We excluded problematic SNPs using standard quality metrics: those with call rate below 95% or Hardy Weinberg P<1<10^-6^ [27]. We also excluded from analysis variants where exome sequencing and Axiom genotyping disagreed on the identity of the major allele and cMAF<1%, because these few variants (32) have extremely large discrepancies in their genotype and are clearly not working in at least one of the two technologies.

### Exome sequencing data

We compared UK Biobank array genotyping data with whole-exome sequencing (WES) data from the UK Biobank to compute summary statistics for samples genotyped in both datasets. We used two releases of WES data from the UK Biobank in multiple formats for this purpose. The first WES data release (50K) consisted of 49,936 individuals [6] that were also processed in the second WES release (200K) along with additional samples for a total of 200,455 samples after exclusion of withdrawn participants [7]. We used the original 50K dataset to replicate the results of Weedon et al. [1] and repeated our analyses with the 200K dataset for a larger sample size.

These two datasets have several important differences in how they were processed. The 50K data was processed with a Functional Equivalence (FE) pipeline that utilized GATK [10] for both per-sample variant calling and joint-genotyping, producing intermediate genomic VCF (gVCF) files for each sample (50K FE-VCF) and a population-level file in PLINK format (50K PLFE-PLINK) after joint-genotyping the 50K FE-VCF files. The 200K data release was processed with a Functional Equivalence protocol that retains original quality scores (OQFE). In contrast to the 50K FE pipeline, the machine-learning based variant caller DeepVariant [11] was used to generate per-sample gVCFs (200K OQFE-VCF) followed by joint-genotyping using GLnexus [12, 15] to produce a population-level VCF file (200K pVCF) that was not filtered prior to release and was the basis for the 200K WES data in PLINK format.

To reproduce the results in Weedon et al., we used the 50K FE-VCF calls with exome sequencing depth>15, as described there [1]. We also analyzed the joint-genotyped 200K OQFE-PLINK data as this release supersedes the 50K data release per UK Biobank and provides a larger sample size. The OQFE-PLINK data uses joint genotyping and is restricted to variants that are polymorphic in the WES data. We used the 200K OQFE-PLINK data without additional quality filters although PPV somewhat improves with stricter quality filters to the 200K OQFE data.

#### Computation and Summary of Performance Metrics

In accordance with previous work, we calculated for each variant four performance metrics: sensitivity, specificity, positive predictive value and negative predictive value. Following Krusche et al’s [5] ‘allele match’ variant matching scheme, we consider any genotype containing the minor allele (i.e., Het or minor Hom) to be a positive and the major Hom to be a negative (we consider only bi-allelic variants in this study). Instances where one of the datasets compared had a No Call were excluded from calculations. For each performance metric we report the number of variants over which the metric is computed. For example, PPV can only be computed for a variant where the array has a Het or minor homozygous call for a sample with WES and sensitivity can only be computed when WES data for the variant has at least one Het or minor homozygous call. We report the median and inter-quartile range of the performance metrics since they are not normally distributed. We also compute the mean PPV for the purpose of comparison with the results of Weedon et al. The focus of this work is the removal of false positives via the RHA algorithm, and this mainly affects the PPV metric. In the very low MAF ranges there is a large excess of true negatives, as a result of which specificity is always extremely high. We therefore report only on PPV in the main text, with data for all four performance metrics reported in the supplementary material.

#### Results and detailed analysis

We directionally reproduced the PPV results obtained by Weedon et al. [1], using first the genomic Variant Call Format (gVCF) files for the 50K exome data restricted to genotype calls covered by more than 15 reads in the sequencing data. Figure 3A and Table 1 show the results on the 50K data before and after the RHA algorithm, showing the significant improvements in PPV for low MAF ranges. In the lowest MAF range (below 0.001%) mean PPVs (computed by taking the PPV for each variant and computing the mean) improved from 16% to 67%, computed over variants with both exome and array data. This lowest MAF range contains less than 5,000 array Het genotypes across only 3,600 variants. In MAF ranges between 0.005%-0.01% PPVs increased to at least 83% while for MAF ranges between 0.01%-1%, PPV increased to 95.5%.

**Fig. 3.**
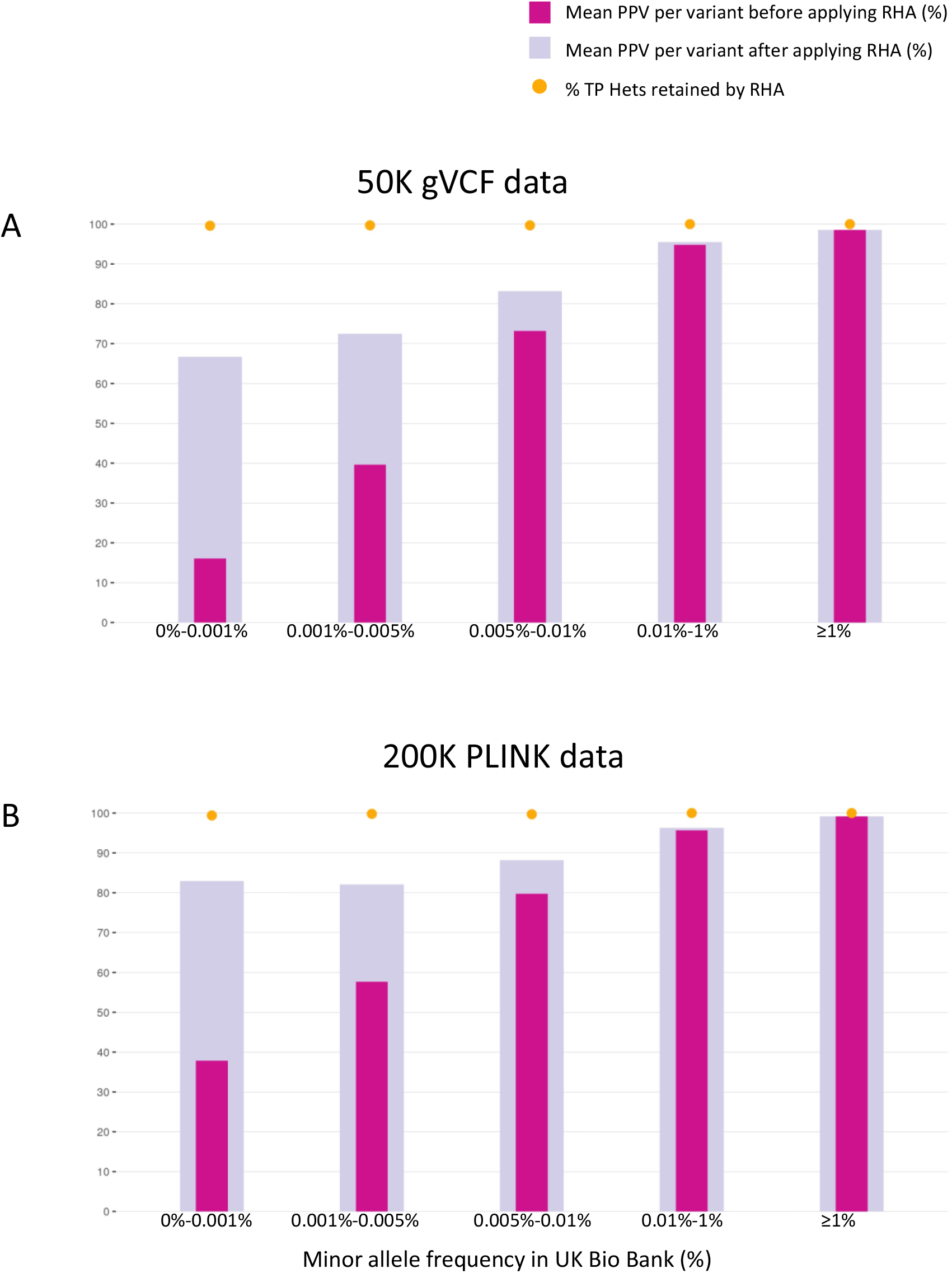
Improvement in positive predictive value of rare variants after application of RHA. Mean positive predictive value (PPV) of variants genotyped by UK BioBank Axiom™ array is shown for different ranges of minor allele frequencies, as calculated from the genotyping results before applying RHA (cMAF). We used genotypes from exome datasets 50K FE-VCF (A) or 200K OQFE-PLINK (B) as truth, comparing performance before and after applying RHA. The percentage of true positive Het calls retained after applying RHA is also indicated. Data for BiLEVE Axiom™ array shows similar trends (Supplementary Material). Note that for a given cMAF range, the number of variants contributing to the mean may be lower after the application of RHA because for some variants RHA eliminates all Het predictions by the array, so that PPV cannot be calculated. Table1 shows the summary statistics of this data, as well as the number of variants contributing to the mean PPV.

We then proceeded to compare array genotyping data before and after RHA to the joint-genotyped 200K OQFE-PLINK data (Figure 3B and Table 1). The 200K OQFE-PLINK data differs from the 50K data in several ways as explained in the previous section. In particular, the 200K joint-genotyped OQFE-PLINK data does not contain loci that are monomorphic in WES (i.e., show no minor alleles in the exome sequencing data).

The difference in PPV (pre-RHA) between Figure 3A and Figure 3B, also seen in Table 1, is largely due to the effect of monomorphic variants in the exome data for which the array data produces a few Hets. The figures also show that post-RHA the difference significantly diminishes, with RHA detecting and removing most false Het calls in these and other variants. The three lowest frequency groups (0,0.001%], (0.001%, 0.005%] and (0.005%, 0.01%] show an initial mean PPV of 38%, 58% and 80% respectively and improve to 83%, 82% and 88%, respectively. Variants with MAF greater than 0.01% but less than 1% and variants with MAF≥1% have an initial mean PPV of 95% and 99%, respectively, and only marginally benefit from the new algorithm.

To illustrate both the content in the various MAF ranges, as well as the large differences in the starting PPV between the 50K gVCF and the 200K PLINK data, we divided the variants in each MAF range into three groups (Fig. 4A, Table 2)

a. Non-responsive in Axiom. Variants that have low MAF in the array data, but have high MAF in the population, based on other evidence. These correspond to non-responsive probesets in the array data. While this group of variants have generally reasonable PPV they have very bad sensitivity and should be filtered from the array data. The number of such variants is small but has significant deleterious effects on the sensitivity of the array data. We defined a variant as non-responsive if the gnomAD [4] MAF of the variant in the Non-Finnish European (NFE) population was significantly higher than the array MAF. Specifically, the variant is considered non-responsive if gnomAD MAF is >= 10*M_r_, where M_r_ equals the upper bound of the array MAF range the variant falls into, i.e. [0, 0.001%), [0.001%, 0.005%), ([0.005%, 0.01%), [0.01%, 1%). Table 3 shows the effect on sensitivity of excluding the non-responsive probesets.
b. Variants that are monomorphic in the exome sequencing data. While this group always has PPV=0 for all variants, even for those with just a single Het in the array data, since by definition of this groups there are no true positive Hets, it also has the largest improvement post-RHA in eliminating spurious Hets. In some sense, while this group can contain very rare variants of interest, the majority of these correspond to loci in the genome that are missing entirely from gnomAD and are likely homozygous in any population. These were included in the array design based on early sequencing results but have proven to be fundamentally not informative in the UK Biobank cohort, since none of the 200K participants for which exome sequencing data is available carry the minor allele. To understand the contribution of these variants to PPV we further break them into 3 subgroups, corresponding to their polymorphic status in the UK Biobank array, (Fig. 4B and Table 4). The 3 subgroups are ‘monomorphic in Axiom pre-RHA’, ‘monomorphic in Axiom only post-RHA’ and ‘polymorphic in Axiom pre- and post-RHA’. The third subgroup is the only one that remains non-concordant with WES post RHA. In the lowest cMAF range (<0.001%), 27% of variants monomorphic in exome do not produce any false positives in the array even without RHA. Applying RHA eliminates almost 90% of false Het predictions in the remaining 73% of these variants, leaving only 970 (13%) of these variants to produce a total of 1,220 Het predictions not in the WES data. For cMAF ranges 0.001% to 0.005% and 0.005% to 0.01%, while a significant proportion of the variants monomorphic in exome remain polymorphic in the array post-RHA (56.1% and 96.2%, respectively), most false positive Hets in this variant class are eliminated by RHA (77.1% and 59.8%, respectively). For cMAF between 0.01% and 1% RHA has mostly little effect on Het predictions because RHA is only applied to small Het clusters, which are rarely found in these MAF ranges.
c. All other variants. These variants are most informative for the computation of PPV and along with Group a are the only ones that contribute to sensitivity.

**Fig. 4.**
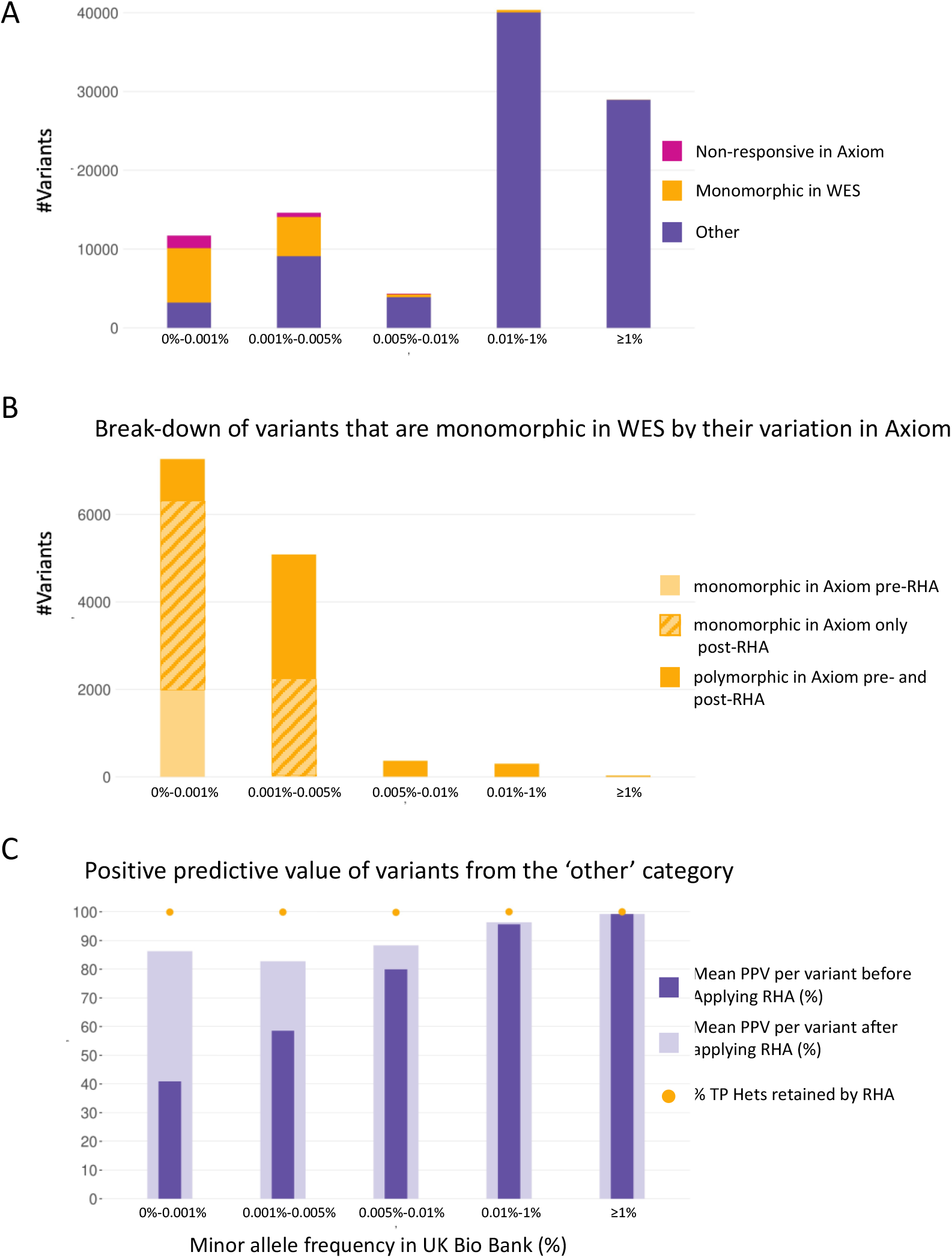
At very low cMAF (≤0.005%) a large proportion of variants are monomorphic in the exome sequencing. A. The number of variants in various cMAF ranges are divided into three groups: monomorphic in the exome sequencing, non-responsive in array data, and other. Note that a variant can be both non-responsive in the array data and monomorphic in the exome sequencing; in this panel such variants were counted in the ‘non-responsive in Axiom’ category. Variants that are monomorphic in exome make up 59% and 34% of the total variants in cMAF ranges 0%-0.001% and 0.001%-0.005%, respectively. B. The variants that are monomorphic in exome sequencing (including those that are also non-responsive in the array data, are further characterized by their variability in Axiom™: monomorphic on the array before applying RHA (on the samples sequenced), monomorphic on the Axiom™ after applying RHA, and polymorphic on Axiom™. C. Mean positive predictive value (PPV) of variants genotyped by UK BioBank Axiom™ Array, using genotypes from 200K OQFE-PLINK as truth, and restricting to variants that are polymorphic in the exome sequencing and responsive in the array data. We show performance before and after applying RHA, as well as the percentage of true positive Het calls retained after applying RHA.

**Table 2.** Distribution of single nucleotide variants with coverage by both Axiom™ UK BioBank array and WES sequencing among three behavior categories. The three categories considered are: ‘non-responsive in Axiom’ (variant’s cMAF is significantly smaller than the non-Finnish European MAF recorded for the variant in gnomAD), ‘monomorphic in WES’ (no minor alleles were found in WES) and ‘other’. Note that for the first three rows the numbers in the three categories do not add up to the total. This is due to a relatively small number of variants (at most 5% of the variants in the cMAF range) that are both ‘non-responsive in Axiom’ and ‘monomorphic in WES’. In Fig. 4A these were counted in the ‘non-responsive in Axiom’ category.

| cMAF range | Total number of variants | Number of variants non-responsive in Axiom | Number of variants monomorphic in WES | Number of 'other' variants |
| --- | --- | --- | --- | --- |
| 0%-0.001% | 11,700 | 1,586 | 7,267 | 3,209 |
| 0.001%-0.005% | 14,619 | 541 | 5,082 | 9,093 |
| 0.005%-0.01% | 4,340 | 84 | 368 | 3,892 |
| 0.01%-1% | 40,350 | 4 | 303 | 40,044 |
| ≥1% | 28,957 | 0 | 29 | 28,928 |

**Table 3.** Effect of exclusion of probesets non-responsive in array on sensitivity. Results are split by the WES dataset used as reference - 50K FE-VCF and 200K OQFE-PLINK, and by the minor allele frequency calculated from the array before application of RHA (cMAF). We include the following summary statistics, calculated across all relevant single nucleotide variants: median and inter-quartile range of sensitivity (the distribution of sensitivity is not normal) as well as the mean sensitivity, for the purposes of comparison with Weedon et al. Also included are the number of variants the summary statistics have been calculated over (only variants with at least one TP or FN genotype can yield a sensitivity value). Insertions and deletions have been excluded from calculations.

| Reference and dataset | Before/<br>After exclusion of<br>non-responsive<br>probesets | Sensitivity (%) |  |  |
| --- | --- | --- | --- | --- |
|  |  | Number<br>of variants | Median (interquartile<br>range) | Mean |
| 50K FE-VCF<br>(n=45,895) |  |  |  |  |
| cMAF 0%-0.001% | Before | 1,999 | 0.0% (0.0%-100.0%) | 28.50% |
|  | After | 1,018 | 100.0% (0.0%-100.0%) | 51.70% |
| cMAF<br>0.001%-0.005% | Before | 6,114 | 100.0% (100.0%-100.0%) | 81.60% |
|  | After | 5,724 | 100.0% (100.0%-100.0%) | 86.50% |
| cMAF<br>0.005%-0.01% | Before | 3,602 | 100.0% (100.0%-100.0%) | 93.00% |
|  | After | 3,533 | 100.0% (100.0%-100.0%) | 94.40% |
| cMAF<br>0.01%-1% | Before | 38,526 | 100.0% (100.0%-100.0%) | 98.40% |
|  | After | 38,537 | 100.0% (100.0%-100.0%) | 98.40% |
| cMAF≥1% | Before | 27,469 | 100.0% (100.0%-100.0%) | 99.90% |
|  | After | 27,469 | 100.0% (100.0%-100.0%) | 99.90% |
| 200K OQFE-PLINK<br>(n=195,447) |  |  |  |  |
| cMAF 0%-0.001% | Before | 3,332 | 33.3% (0.0%-100.0%) | 49.50% |
|  | After | 2,216 | 100.0% (0.0%-100.0%) | 70.00% |
| cMAF<br>0.001%-0.005% | Before | 8,757 | 100.0% (100.0%-100.0%) | 84.10% |
|  | After | 8,329 | 100.0% (100.0%-100.0%) | 87.80% |
| cMAF<br>0.005%-0.01% | Before | 3,943 | 100.0% (100.0%-100.0%) | 92.80% |
|  | After | 3,864 | 100.0% (100.0%-100.0%) | 94.30% |
| cMAF<br>0.01%-1% | Before | 40,009 | 100.0% (100.0%-100.0%) | 98.40% |
|  | After | 40,006 | 100.0% (100.0%-100.0%) | 98.40% |
| cMAF<br>≥1% | Before | 28,893 | 100.0% (100.0%-100.0%) | 99.90% |
|  | After | 28,893 | 100.0% (100.0%-100.0%) | 99.90% |

**Table 4.**
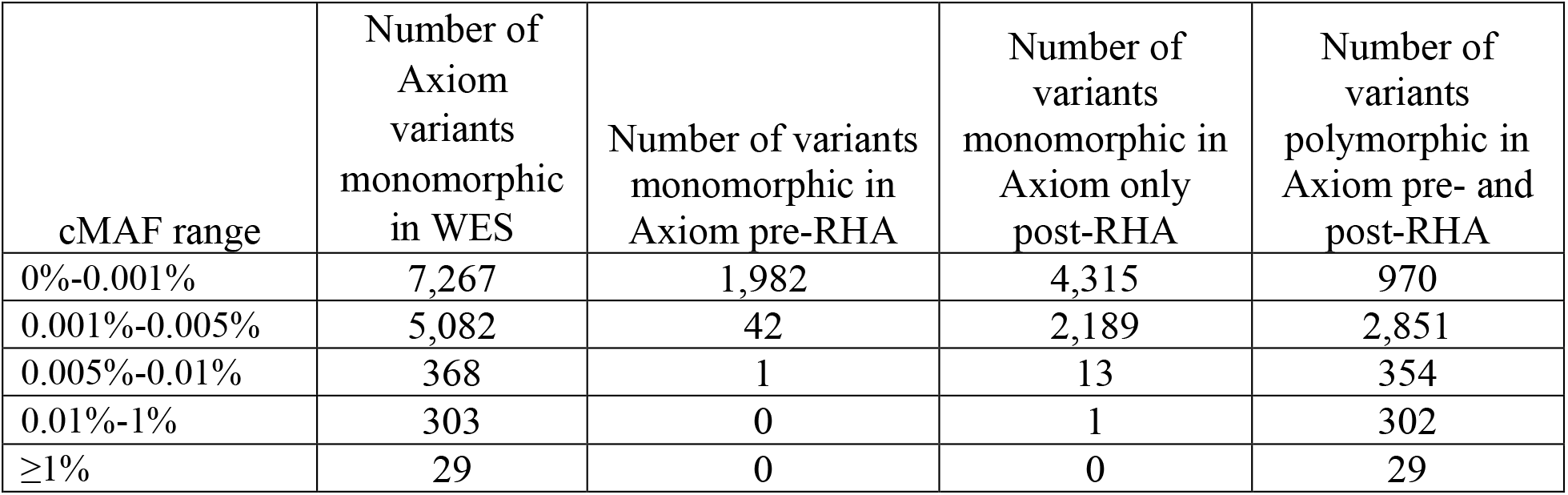
Polymorphic status in Axiom™ UKB BioBank array of single nucleotide variants that are monomorphic in 200K OQFE-PLINK WES. Variants are divided into three categories: ‘monomorphic in Axiom pre-RHA’, ‘monomorphic in Axiom only post-RHA’ and polymorphic in Axiom pre- and post-RHA. Note that for the purpose of this classification we consider for each variant only samples that have a genotype in both Axiom and WES, so that a variant can be both monomorphic in Axiom pre-RHA and have cMAF>0.

| cMAF range | Number of Axiom variants monomorphic in WES | Number of variants monomorphic in Axiom pre-RHA | Number of variants monomorphic in Axiom only post-RHA | Number of variants polymorphic in Axiom pre- and post-RHA |
| --- | --- | --- | --- | --- |
| 0%-0.001% | 7,267 | 1,982 | 4,315 | 970 |
| 0.001%-0.005% | 5,082 | 42 | 2,189 | 2,851 |
| 0.005%-0.01% | 368 | 1 | 13 | 354 |
| 0.01%-1% | 303 | 0 | 1 | 302 |
| ≥1% | 29 | 0 | 0 | 29 |

Group a (non-responsive variants) constitutes 13.6% of variants with cMAF<0.001% but has few variants in higher allele frequency ranges, as shown in Figure 4a.

Group b (monomorphic in WES) has a large number of variants pre RHA in the two lowest frequency groups, but most of these are eliminated post RHA as shown in Figure 4b. As expected, Group b has almost no variants with array frequencies above 0.01%. Group b is not represented in the 200K PLINK data, as mentioned earlier.

#### Changes in PPV resulting from increased QC of exome sequencing

As the 200K WES data was not filtered prior to release [17], we tested a range of cutoff values for sequencing depth and genotype quality score to determine the effect of stronger quality control on our results. Sequencing depth appeared to be a factor, but we noticed that the sequencing depth of major homozygous calls in the exome data is significantly lower than the sequencing depth for exome Het calls, Figure 5. We therefore abandoned the attempt to filter exome calls by sequencing depth to avoid introducing an even larger bias.

**Fig. 5.**
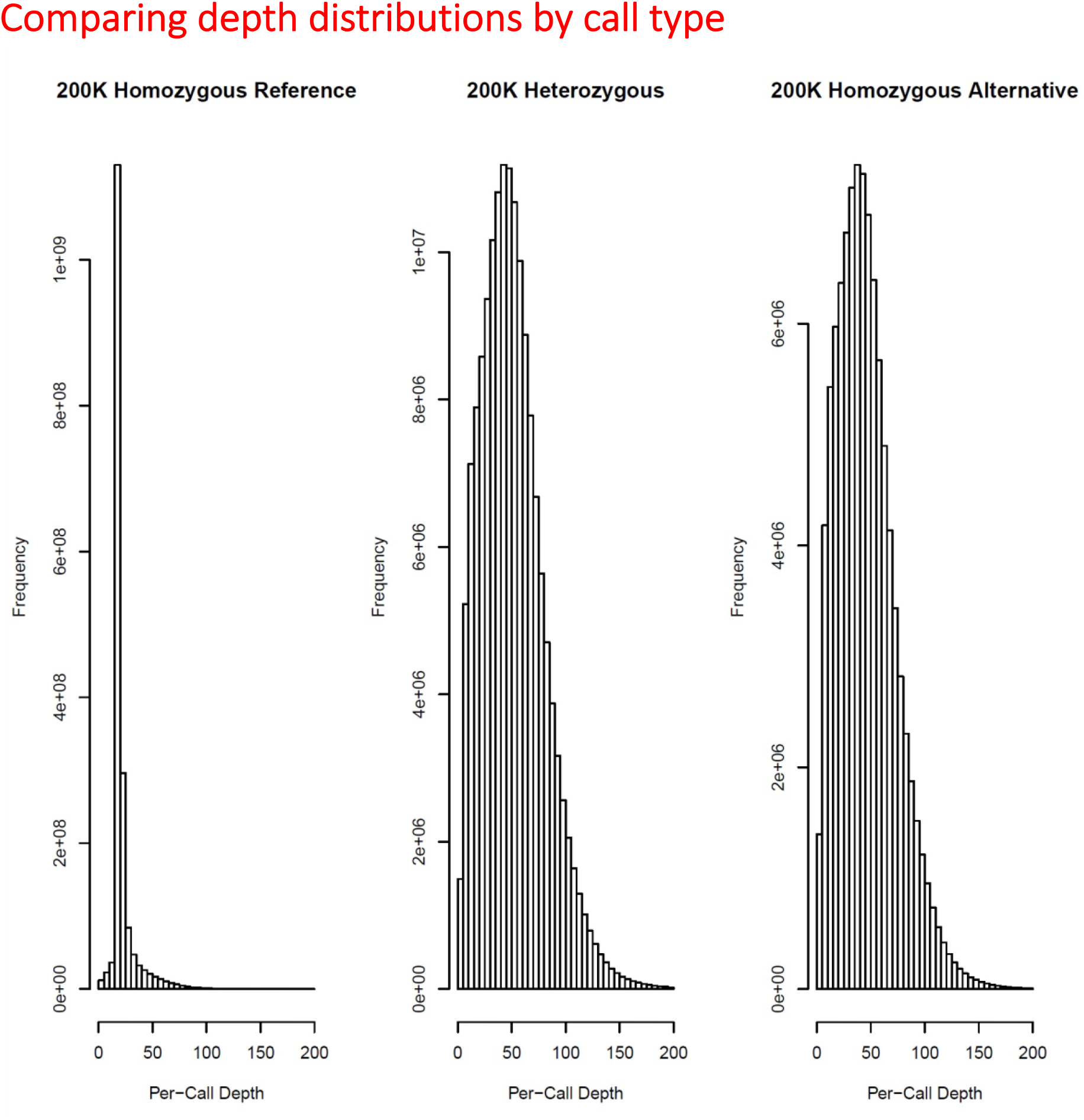
Comparison of WES 200K sequencing depth by exome genotyping call. We show that the depth of exome sequencing by genotyping call, homozygous ref, het or homozygous alt. Sequencing depth for exome homozygous ref calls have significantly lower sequencing depth than exome het calls. Quality filtering by depth would therefore disproportionally filter out homozygous ref calls.

#### Probeset Filtering

Probesets can be filtered in many ways, most notably by pre-qualifying probesets based on genotyping results for known samples. In the current analysis the only filtering we applied was the most obvious one, eliminating non-responsive probesets, where the known MAF is significantly higher than the computed MAF in the array data (see Table 5).

**Table 5.** Positive predictive value of UK BioBank Axiom™ array versus whole exome sequencing before and after application of RHA - effect of adding/removing variants monomorphic in WES or non-responsive in Axiom. In both parts of the table 200K OQFE-PLINK is used as a reference and results are split by the minor allele frequency calculated from the array before application of RHA (cMAF). In the first part of the table, we add variants monomorphic in 50K FE-VCF but absent from 200K OQFE-PLINK to the analysis of variants in 200K OQFE-PLINK. For these added variants, all positive predictions in the samples sequenced by WES are assumed to be false positives and they can therefore only contribute PPV values of 0 to the distribution (or be excluded from the distribution if they have no positive predictions). In the second part of the table, we exclude from the comparison variants non-responsive in Axiom or monomorphic in WES. For each dataset, we include the following summary statistics, calculated across all relevant single nucleotide variants: median and inter-quartile range of PPV (the distribution of PPV is not normal) as well as the mean PPV, for the purposes of comparison with Weedon et al. Also included are the number of variants the summary statistics have been calculated over (only variants with at least one TP or FP genotype can yield a PPV) and the percentage of TP Hets that are retained after applying RHA measured as (TP hets pre RHA)/(TP hets post RHA) (only Het genotypes are affected by the algorithm). Insertions and deletions have been excluded from calculations.

| Reference and dataset | Before/<br>After/<br>application of<br>RHA | Positive Predictive Value (%) |  |  | %TP Hets<br>retained<br>after RHA |
| --- | --- | --- | --- | --- | --- |
|  |  | Number of<br>variants | Median (interquartile<br>range) | Mean |  |
| 200K OQFE-PLINK<br>(n=195,447), adding<br>variants monomorphic<br>in WES |  |  |  |  |  |
| All exome variants | Before | 96,620 | 99.0% (85.7%- 99.9%) | 79.5% |  |
|  | After | 88,157 | 99.6% (97.4%-100.0%) | 90.3% | >99.9% |
| cMAF 0%-0.001% | Before | 8,646 | 0.0% (0.0%- 0.0%) | 14.7% |  |
|  | After | 3,104 | 100.0% (0.0%-100.0%) | 57.1% | 99.5% |
| cMAF<br>0.001%-0.005% | Before | 14,397 | 20.0% (0.0%- 80.0%) | 37.5% |  |
|  | After | 11,495 | 90.9% (0.0%-100.0%) | 61.7% | 99.8% |
| cMAF<br>0.005%-0.01% | Before | 4,328 | 89.0% (61.1%- 97.2%) | 73.0% |  |
|  | After | 4,310 | 97.0% (80.0%-100.0%) | 80.9% | 99.7% |
| cMAF<br>0.01%-1% | Before | 40,323 | 98.9% (96.4%- 99.8%) | 95.0% |  |
|  | After | 40,322 | 99.1% (97.0%- 99.8%) | 95.6% | >99.9% |
| cMAF≥1% | Before | 28,926 | 99.9% (99.6%-100.0%) | 99.1% | |
|  | After | 28,926 | 99.9% (99.6%-100.0%) | 99.1% | 100.0% |
| 200K OQFE-PLINK<br>(n=195,447),<br>removing variants<br>non-responsive in<br>Axiom or<br>monomorphic in WES |  |  |  |  |  |
| All exome variants | Before | 84,360 | 99.4% (95.9%- 99.9%) | 90.5% |  |
|  | After | 82,867 | 99.7% (98.1%-100.0%) | 95.4% | >99.9% |
| cMAF 0%-0.001% | Before | 2,637 | 33.3% (0.0%- 75.0%) | 40.9% |  |
|  | After | 1,782 | 100.0% (100.0%-100.0%) | 86.3% | 99.9% |
| cMAF<br>0.001%-0.005% | Before | 8,927 | 66.7% (25.0%- 93.3%) | 58.5% |  |
|  | After | 8,294 | 100.0% (80.0%-100.0%) | 82.8% | 99.9% |
| cMAF<br>0.005%-0.01% | Before | 3,882 | 91.3% (73.1%-100.0%) | 80.0% |  |
|  | After | 3,877 | 100.0% (88.9%-100.0%) | 88.3% | 99.8% |
| cMAF<br>0.01%-1% | Before | 40,017 | 98.9% (96.5%- 99.8%) | 95.7% |  |
|  | After | 40,017 | 99.1% (97.1%- 99.9%) | 96.3% | >99.9% |
| cMAF<br>≥1% | Before | 28,897 | 99.9% (99.6%-100.0%) | 99.2% | |
|  | After | 28,897 | 99.9% (99.6%-100.0%) | 99.2% | 100.0% |

#### Variants in the BRCA genes

Weedon et al. [1] identified 1139 variants of interest in *BRCA1* and *BRCA2*. After manual QC of WES data, they found that 425 of these variants had at least one array Het call as well as WES data. Of these 425 variants, only 53 are present on the UK Biobank array; the remaining 372 are only on the BiLEVE array, which was used to genotype the first 10% of UK Biobank participants, comprising only 8% of the 50K WES cohort, and only 3% of the 200K WES cohort.

Weedon et al. [1] also identified 65 individuals with one FN each among the UK Biobank participants, but 60 of these FNs were BiLEVE-only variants that were not actually assayed on the UK Biobank array. This was the result of Weedon et al. assuming all variants were present on both arrays. When considering only variants that are present on the UK Biobank array, and only participants with genotypes from that array, we find 37 exome Het calls in the 50K WES cohort; 32 of these exome Het calls were also called Het on the UK Biobank array. Of the 5 extra WES Het calls (across 5 individuals) 3 received an array ‘No Call’, leaving 2 FNs for an overall sensitivity of 94.1%. The remaining 60 individuals did indeed have a WES Het call in one of the BRCA variants of interest, but as mentioned earlier, it was on a variant not present on the UK Biobank array. The large number of BRCA variants only present on the BiLEVE array, emphasizes the exploratory nature of the probesets for these BRCA variants.

Both arrays did indeed have a significant number of False Positives (FPs). In the 50K WES cohort, the 53 BRCA variants with array positives on the UK Biobank array yielded a total of 189 array Hets, of which 157 were not reproduced by WES. While 16 of these 157 were no-calls in the 50K WES data, the corresponding variants are monomorphic in the 200K WES data, and we therefore added them to the FP count. Thus, in agreement with Weedon et al, the overall pre RHA PPV is 16.9%. RHA eliminated 112 (71.3%) of these FPs, while preserving all true positives, thus improving PPV to 41.6%.

In total among the subset of the 1139 variants of interest, only 91 are on UKBiobank and 1089 are on UKBiLEVE.

We further note that the UK Biobank array has 460 BRCA variants with cMAF < 0.01%, that are included in the 200K WES PLINK data set. Pre RHA these include 4740 positives called by the array, 2983 positives called by Exome Sequencing and 2529 called by both, for a Pre RHA sensitivity of 84.8% and PPV of 53.4%. RHA removed 1613 (73%) of the false positives and 3 (0.1%) true positives, increasing the overall PPV to 80.9% while reducing the sensitivity to 84.7%.

Table 6 and Supplementary Table F contain additional summary statistics for BRCA performance on the UK Biobank and BiLEVE arrays.

**Table 6.** Performance of Axiom™ UK Biobank array on the BRCA module. WES data from 50K WES (left) or 200K OQFE-PLINK (right) is used as a reference. Within each sub-table, the variants included were those studied by Weedon et al [1]. Of the 91 variants present on the UK Biobank array, 54 variants were included in the list of variants with array hets or Exome hets that Weedon used in their analysis and shared with us. The 200K WES release followed these same variants but excludes variants that are monomorphic and hence missing in the 200K WES exome PLINK data. Each data point represents one pair of genotyping calls on one sample. For the 50K WES release, all Array Het, Exome No-Call pairs were manually reviewed and assigned as True Positive or False Negative; for the 200K WES release, instances where one of the two technologies was a no-call were excluded from the counts.

| 50K Exome data (gVCF) |  |  | 200K Exome data (PLINK) |  |
| --- | --- | --- | --- | --- |
| Variants | N = 91 |  | Variants | N = 26 |
| True Positive | 32 |  | True Positive | 141 |
| False Positive | 157 |  | False Positive | 199 |
| False Negative | 2 |  | False Negative | 12 |
| Exome Het<br>Array NC | 3 |  | Exome Het<br>Array NC | 21 |
| Sensitivity | 94.1% |  | Sensitivity | 92.2% |
| PPV (Pre-RHA) | 16.9% |  | PPV (Pre-RHA) | 41.5% |
| PPV (Post-RHA) | 41.6% |  | PPV (Post-RHA) | 65.3% |

## Discussion

Very rare variants are increasingly included on genotyping microarrays [19 – 23]. Since microarrays are not generally suitable for discovery of novel polymorphisms, these rare variants are mostly known from the literature or public databases [24, 25], and have previously established or suspected significance. Rare variants are frequently included on arrays for research purposes, which are indeed Research use only, and raw results should not be returned to naïve test subjects or customers of direct-to-consumer genetics companies [26]. Relevant to the main research use of these arrays, statistical power to detect associations to very rare variants in non-case-control studies is limited. Sporadic false positives or negatives, at a low rate that would leave the power of more common variants essentially unchanged, could have a much greater effect on the analysis of very rare variants.

Weedon et al., using the recently released whole exome sequencing data for 50K UK Biobank participants, observed that rare variants in the UK Biobank data decreased in accuracy as a rough function of minor allele frequency (as computed from the array-derived genotype calls). Variants in the BRCA1 and BRCA2 genes were used as an illustrative example and while sensitivity on these variants was high, PPV before the introduction of the rare het adjusted (RHA) genotyping was low. RHA had been in development and the Biobank array-derived genotype calls along with the WES data for 50K participants provided an excellent test case and greatly decreased the number of false positive heterozygote calls for rare variants without affecting true positive calls. We also used the power of multiple, large sequencing projects, collected in the Genome Aggregation Database (gnomAD) [4], to filter variants by expected allele frequencies – an operation that was less possible and less accurate at the time of the original calling.

We have shown that microarrays can be used for correctly genotyping very rare variants provided that

1. the probeset used for genotyping has been shown to be responsive to the minor allele and has been carefully designed.
2. improved algorithms are used to eliminate spurious heterozygous calls.

For the variants in the BRCA genes, we have shown that a large number of variants perform well, with overall high sensitivity, but a few poorly performing probesets can significantly affect the overall PPV. In addition, a large number of poor performing variants in the BRCA genes were only present on the BiLEVE array (used to genotype the first 50K individuals). That array also provided opportunity for additional quality control. Indeed, many of these variants were subsequently excluded from the UK Biobank design.

Weedon et al. state that because genotyping arrays typically assay many thousands of rare variants simultaneously, and have a specificity that is less than 100%, false positive results will occur and outnumber true positives across all rare variants [1]. This no longer holds after the introduction of the new RHA genotyping algorithm which show a mean PPV above 80% in all MAF ranges and near or above 90% in MAF ranges above 0.005%, and a median PPV near 100% for all MAF ranges.

## Conclusion

The Axiom UK Biobank array is a research use only array and has provided the community with a large number of insightful, unique and consequential research results (https://www.ukbiobank.ac.uk/enable-your-research/publications). The new calling algorithm and probeset filtering described here further and significantly improve the accuracy of the resulting data, show high concordance to WES sequencing data, and provide an important resource for the research community. Weedon et al. also conclude that clinicians should validate array results from direct-to-consumer companies or research biobanks by using a standard diagnostic test before recommending any action. These recommendations apply and should be standard practice for all RUO products, including the WES Biobank data.

## Glossary

*Allele*: each of two or more alternative forms of DNA that are found at the same location on a chromosome
*Allele A and allele B*: For a SNP the two alternatives that can be observed and measured in a given sample are designated as “allele A” and “allele B”
*Array*: DNA microarray that is used to genotype known genetic variants (SNPs and indels) in the population
*Clustering space*: *The X and Y dimensions defined by Signal Contrast and Signal Size*
*Exome*: ~1-2% of the human genome that codes for proteins
*Genotyping*: method for determining the base (A, G, T, or C) present at a specific location in a person’s DNA
*Het call*: short for heterozygous call
*Hom call*: short for homozygous call
*Heterozygous*: two different alleles at a given locus in an individual
*Homozygous*: two identical alleles at a given locus in an individual
*Indel*: type of variant where one or more bases are inserted or deleted as compared to the reference genome
*Major Homozygous*: two identical alleles at a given locus that represent the most common allele at a given locus for the population of interest
*Minor Homozygous*: two identical alleles at a given locus that represent the less common allele at a given locus for the population of interest
*Mean positive predictive value over a set of variants and individuals*: the average over the positive predictive values for each variant
nAB: the number of samples called heterozygous by AxiomGT1 in the clustering space
*Negative predictive value*: proportion of the normal alleles that are confirmed by the reference standard (true negative/(true negative + false negative))
*Positive predictive value*: proportion of variant alleles that are confirmed by the reference standard (true positive/(true positive + false positive))
*Overall positive predictive value*: Positive predictive value across all variants and samples
*Probeset*: A specific set of DNA sequences on the microarray that detect the presence of two or more alleles at a given locus
*Sensitivity*: proportion of variant alleles detected by the reference standard that are also found by the index test (true positive/(true positive + false negative))
*Signal Contrast and Signal Size*: AxiomGT1 genotype clustering is carried out in two dimensions. The X dimension is called “contrast” and the Y dimension is called “size”. They are log-linear combinations of the two allele signal intensities. For alleles A and B, contrast is log2(A/B) and size is (log2(A) +log2(B))/2
*Single nucleotide polymorphism* (SNP): type of single nucleotide variant; a position in the genome where an individual differs from the reference human genome by a single base change (i.e., a substitution of a single letter of DNA). A SNP may be rare or common in the population
*Specificity*: proportion of normal alleles detected by the reference standard that are also found to be normal by the index test (true negative/(true negative + false positive))
*Variant*: Locus in the genome where different alleles have been observed in different people

## Acknowledgments

Analysis of UK Biobank data is conducted using UK Biobank Resource under Application Number 55681. We thank Mike Weedon and Caroline Wright for sharing data with us that was mentioned but not included in their manuscript. [1]

## Notes

### Competing Interest Statement

The authors are employed by Thermo Fisher Scientific, which participated in the design of and manufactures the UK Biobank Axiom Array.

