## Supplemental Tables for "Novel genotyping algorithms for rare variants significantly improve the accuracy of Applied Biosystems ™ Axiom ™ array genotyping calls"

### Supplementary Table A

| Reference and dataset | Before/ After application of RHA | Positive Predictive value (%) | | | Negative Predictive Value (%) | | | Sensitivity (%) | | | Specificity (%) | | | %TP Hets retained after RHA |
|---|---|---|---|---|---|---|---|---|---|---|---|---|---|---|
| | | Number of variants | Median (interquartile range) | Mean | Number of variants | Median (interquartile range) | Mean | Number of variants | Median (interquartile range) | Mean | Number of variants | Median (interquartile range) | Mean | |
| 50K FE-VCF (n=45,895) | | | | | | | | | | | | | | |
| All exome variants | Before | 85,447 | 99.7% (95.0%-100.0%) | 84.5% | 96,813 | 100.0% (100.0%-100.0%) | 100.0% | 77,714 | 100.0% (100.0%-100.0%) | 95.6% | 96,813 | 100.0% (100.0%-100.0%) | 99.9% | |
| | After | 78,468 | 99.9% (98.4%-100.0%) | 93.6% | 96,813 | 100.0% (100.0%-100.0%) | 100.0% | 77,713 | 100.0% (100.0%-100.0%) | 95.6% | 96,813 | 100.0% (100.0%-100.0%) | 99.9% | >99.9% |
| cMAF 0%-0.001% | Before | 3,568 | 0.0% (0.0%-0.0%) | 16.0% | 11,533 | 100.0% (100.0%-100.0%) | 99.9% | 1,999 | 0.0% (0.0%-100.0%) | 28.5% | 11,533 | 100.0% (100.0%-100.0%) | 100.0% | |
| | After | 934 | 100.0% (0.0%-100.0%) | 66.9% | 11,533 | 100.0% (100.0%-100.0%) | 99.9% | 1,999 | 0.0% (0.0%-100.0%) | 28.5% | 11,533 | 100.0% (100.0%-100.0%) | 100.0% | 99.6% |
| cMAF 0.001%-0.005% | Before | 11,134 | 0.0% (0.0%-100.0%) | 39.6% | 14,266 | 100.0% (100.0%-100.0%) | 99.9% | 6,115 | 100.0% (100.0%-100.0%) | 81.6% | 14,266 | 100.0% (100.0%-100.0%) | 100.0% | |
| | After | 6,968 | 100.0% (50.0%-100.0%) | 72.6% | 14,266 | 100.0% (100.0%-100.0%) | 99.9% | 6,114 | 100.0% (100.0%-100.0%) | 81.6% | 14,266 | 100.0% (100.0%-100.0%) | 100.0% | 99.7% |
| cMAF 0.005%-0.01% | Before | 4,107 | 90.9% (57.1%-100.0%) | 73.2% | 4,188 | 100.0% (100.0%-100.0%) | 99.9% | 3,602 | 100.0% (100.0%-100.0%) | 93.0% | 4,188 | 100.0% (100.0%-100.0%) | 100.0% | |
| | After | 3,954 | 100.0% (83.3%-100.0%) | 83.2% | 4,188 | 100.0% (100.0%-100.0%) | 99.9% | 3,602 | 100.0% (100.0%-100.0%) | 93.0% | 4,188 | 100.0% (100.0%-100.0%) | 100.0% | 99.7% |
| cMAF 0.01%-1% | Before | 38,980 | 99.2% (96.3%-100.0%) | 94.8% | 39,107 | 100.0% (100.0%-100.0%) | 100.0% | 38,529 | 100.0% (100.0%-100.0%) | 98.4% | 39,107 | 100.0% (100.0%-100.0%) | 100.0% | |
| | After | 38,954 | 99.4% (96.9%-100.0%) | 95.5% | 39,107 | 100.0% (100.0%-100.0%) | 100.0% | 38,529 | 100.0% (100.0%-100.0%) | 98.4% | 39,107 | 100.0% (100.0%-100.0%) | 100.0% | >99.9% |
| cMAF≥1% | Before | 27,658 | 99.9% (99.6%-100.0%) | 98.6% | 27,719 | 100.0% (100.0%-100.0%) | 100.0% | 27,469 | 100.0% (100.0%-100.0%) | 99.9% | 27,719 | 100.0% (99.9%-100.0%) | 99.5% | |
| | After | 27,658 | 99.9% (99.6%-100.0%) | 98.6% | 27,719 | 100.0% (100.0%-100.0%) | 100.0% | 27,469 | 100.0% (100.0%-100.0%) | 99.9% | 27,719 | 100.0% (99.9%-100.0%) | 99.5% | 100.0% |
| 200K OQFE-PLINK (n=195,447) | | | | | | | | | | | | | | |
| All exome variants | Before | 85,598 | 99.4% (95.6%-99.9%) | 89.7% | 86,501 | 100.0% (100.0%-100.0%) | 100.0% | 84,936 | 100.0% (100.0%-100.0%) | 95.3% | 86,501 | 100.0% (100.0%-100.0%) | 99.9% | |
| | After | 83,652 | 99.7% (98.1%-100.0%) | 95.1% | 86,501 | 100.0% (100.0%-100.0%) | 100.0% | 84,934 | 100.0% (100.0%-100.0%) | 95.2% | 86,501 | 100.0% (100.0%-100.0%) | 99.9% | >99.9% |

| Reference and dataset | Before/ After application of RHA | Positive Predictive value (%) | | | Negative Predictive Value (%) | | | Sensitivity (%) | | | Specificity (%) | | | %TP Hets retained after RHA |
|---|---|---|---|---|---|---|---|---|---|---|---|---|---|---|
| | | Number of variants | Median (interquartile range) | Mean | Number of variants | Median (interquartile range) | Mean | Number of variants | Median (interquartile range) | Mean | Number of variants | Median (interquartile range) | Mean | |
| cMAF 0%-0.001% | Before | 3,361 | 33.3% (0.0%-66.7%) | 37.8% | 4,235 | 100.0% (100.0%-100.0%) | 99.7% | 3,333 | 33.3% (0.0%-100.0%) | 49.5% | 4,235 | 100.0% (100.0%-100.0%) | 100.0% | |
| | After | 2,134 | 100.0% (100.0%-100.0%) | 83.0% | 4,235 | 100.0% (100.0%-100.0%) | 99.7% | 3,332 | 33.3% (0.0%-100.0%) | 49.5% | 4,235 | 100.0% (100.0%-100.0%) | 100.0% | 99.5% |
| cMAF 0.001%-0.005% | Before | 9,358 | 66.7% (22.2%-93.3%) | 57.7% | 9,385 | 100.0% (100.0%-100.0%) | 99.9% | 8,757 | 100.0% (100.0%-100.0%) | 84.1% | 9,385 | 100.0% (100.0%-100.0%) | 100.0% | |
| | After | 8,644 | 100.0% (75.0%-100.0%) | 82.1% | 9,385 | 100.0% (100.0%-100.0%) | 99.9% | 8,757 | 100.0% (100.0%-100.0%) | 84.1% | 9,385 | 100.0% (100.0%-100.0%) | 100.0% | 99.8% |
| cMAF 0.005%-0.01% | Before | 3,961 | 91.3% (72.7%-100.0%) | 79.8% | 3,962 | 100.0% (100.0%-100.0%) | 100.0% | 3,944 | 100.0% (100.0%-100.0%) | 92.9% | 3,962 | 100.0% (100.0%-100.0%) | 100.0% | |
| | After | 3,956 | 100.0% (88.9%-100.0%) | 88.1% | 3,962 | 100.0% (100.0%-100.0%) | 100.0% | 3,943 | 100.0% (100.0%-100.0%) | 92.8% | 3,962 | 100.0% (100.0%-100.0%) | 100.0% | 99.7% |
| cMAF 0.01%-1% | Before | 40,021 | 98.9% (96.5%-99.8%) | 95.7% | 40,022 | 100.0% (100.0%-100.0%) | 100.0% | 40,009 | 100.0% (100.0%-100.0%) | 98.4% | 40,022 | 100.0% (100.0%-100.0%) | 100.0% | |
| | After | 40,021 | 99.1% (97.1%-99.9%) | 96.3% | 40,022 | 100.0% (100.0%-100.0%) | 100.0% | 40,009 | 100.0% (100.0%-100.0%) | 98.4% | 40,022 | 100.0% (100.0%-100.0%) | 100.0% | >99.9% |
| cMAF≥1% | Before | 28,897 | 99.9% (99.6%-100.0%) | 99.2% | 28,897 | 100.0% (100.0%-100.0%) | 100.0% | 28,893 | 100.0% (100.0%-100.0%) | 99.9% | 28,897 | 100.0% (99.9%-100.0%) | 99.7% | |
| | After | 28,897 | 99.9% (99.6%-100.0%) | 99.2% | 28,897 | 100.0% (100.0%-100.0%) | 100.0% | 28,893 | 100.0% (100.0%-100.0%) | 99.9% | 28,897 | 100.0% (99.9%-100.0%) | 99.7% | 100.0% |

**Performance of UK BioBank Axiom™ array versus whole exome sequencing before and after application of RHA.** Results are split by the WES dataset used as reference -50K FE-VCF and 200K OQFE-PLINK, and by the minor allele frequency calculated from the array before application of RHA (cMAF). We include the following performance metrics: positive predictive value (PPV), negative predictive value (NPV), sensitivity and specificity, as well as the percentage of TP Hets that are retained after applying RHA (only Het genotypes are affected by the algorithm). For each performance metric, the following summary statistics are provided, calculated across all relevant single nucleotide variants: median and inter-quartile range of PPV (the distribution of PPV is not normal) as well as the mean PPV, for the purposes of comparison with Weedon et al. Also included are the number of variants for which the performance metric could be calculated. Variants corresponding to insertions and deletions have been excluded from calculations.

### Supplementary Table B

| Reference and dataset | Before/after application of RHA | Number of variants with positive predictive value | Overall positive predictive value |
|---|---|---|---|
| 50K FE-VCF (n=45,895) | | | |
| All exome variants | Before | 85,479 | 98.9% |
| | After | 78,500 | 98.9% |
| cMAF 0%-0.001% | Before | 3,574 | 16.5% |
| | After | 940 | 67.9% |
| cMAF 0.001%-0.005% | Before | 11,147 | 45.0% |
| | After | 6,981 | 79.3% |
| cMAF 0.005%-0.01% | Before | 4,107 | 74.5% |
| | After | 3,954 | 88.3% |
| cMAF 0.01%-1% | Before | 38,993 | 97.9% |
| | After | 38,967 | 98.0% |
| cMAF≥1% | Before | 27,658 | 98.9% |
| | After | 27,658 | 98.9% |
| 200K OQFE-PLINK (n=195,447) | | | |
| All exome variants | Before | 85,621 | 99.2% |
| | After | 83,675 | 99.2% |
| cMAF 0%-0.001% | Before | 3,365 | 38.6% |
| | After | 2,138 | 85.1% |
| cMAF 0.001%-0.005% | Before | 9,365 | 62.4% |
| | After | 8,651 | 88.4% |
| cMAF 0.005%-0.01% | Before | 3,961 | 80.6% |
| | After | 3,956 | 91.3% |
| cMAF 0.01%-1% | Before | 40,033 | 98.0% |
| | After | 40,033 | 98.1% |
| cMAF≥1% | Before | 28,897 | 99.3% |
| | After | 28,897 | 99.3% |

Overall positive predictive value of UK BioBank Axiom™ array versus whole exome sequencing before and after application of RHA. Results are split by the WES dataset used as reference -50K FE-VCF and 200K OQFE-PLINK, and by the minor allele frequency calculated from the array before application of RHA (cMAF). Also included are the number of variants for which positive predictive value could be calculated. Variants corresponding to insertions and deletions have been excluded from calculations.

### Supplementary Table C

| Reference and dataset | Before/ After application of RHA | Positive Predictive value (%) | | | Negative Predictive Value (%) | | | Sensitivity (%) | | | Specificity (%) | | | %TP Hets retained after RHA |
|---|---|---|---|---|---|---|---|---|---|---|---|---|---|---|
| | | Number of variants | Median (interquartile range) | Mean | Number of variants | Median (interquartile range) | Mean | Number of variants | Median (interquartile range) | Mean | Number of variants | Median (interquartile range) | Mean | |
| 50K FE-VCF (n=4,041) | | | | | | | | | | | | | | |
| All exome variants | Before | 65,569 | 100.0% (99.1%-100.0%) | 91.0% | 86,040 | 100.0% (100.0%-100.0%) | 100.0% | 63,121 | 100.0% (100.0%-100.0%) | 96.4% | 86,041 | 100.0% (100.0%-100.0%) | 99.8% | |
| | After | 62,374 | 100.0% (99.6%-100.0%) | 96.0% | 86,040 | 100.0% (100.0%-100.0%) | 100.0% | 63,118 | 100.0% (100.0%-100.0%) | 96.4% | 86,041 | 100.0% (100.0%-100.0%) | 99.8% | >99.9% |
| cMAF 0%-0.001% | Before | 524 | 0.0% (0.0%-0.0%) | 8.6% | 7,772 | 100.0% (100.0%-100.0%) | 99.8% | 849 | 0.0% (0.0%-0.0%) | 4.8% | 7,772 | 100.0% (100.0%-100.0%) | 100.0% | |
| | After | 79 | 100.0% (0.0%-100.0%) | 57.0% | 7,772 | 100.0% (100.0%-100.0%) | 99.8% | 849 | 0.0% (0.0%-0.0%) | 4.8% | 7,772 | 100.0% (100.0%-100.0%) | 100.0% | 97.80% |
| cMAF 0.001%-0.005% | Before | 2,711 | 0.0% (0.0%-0.0%) | 18.3% | 10,429 | 100.0% (100.0%-100.0%) | 99.9% | 1,011 | 13.3% (0.0%-100.0%) | 48.9% | 10,429 | 100.0% (100.0%-100.0%) | 100.0% | |
| | After | 784 | 100.0% (0.0%-100.0%) | 65.5% | 10,429 | 100.0% (100.0%-100.0%) | 99.9% | 1,011 | 13.3% (0.0%-100.0%) | 48.9% | 10,429 | 100.0% (100.0%-100.0%) | 100.0% | 99.70% |
| cMAF 0.005%-0.01% | Before | 1,183 | 100.0% (0.0%-100.0%) | 58.7% | 3,152 | 100.0% (100.0%-100.0%) | 99.9% | 857 | 100.0% (100.0%-100.0%) | 83.4% | 3,152 | 100.0% (100.0%-100.0%) | 100.0% | |
| | After | 836 | 100.0% (100.0%-100.0%) | 85.9% | 3,152 | 100.0% (100.0%-100.0%) | 99.9% | 857 | 100.0% (100.0%-100.0%) | 83.4% | 3,152 | 100.0% (100.0%-100.0%) | 100.0% | 100.00% |
| cMAF 0.01%-1% | Before | 33,856 | 100.0% (98.0%-100.0%) | 93.2% | 37,269 | 100.0% (100.0%-100.0%) | 100.0% | 33,305 | 100.0% (100.0%-100.0%) | 97.6% | 37,269 | 100.0% (100.0%-100.0%) | 100.0% | |
| | After | 33,380 | 100.0% (100.0%-100.0%) | 94.9% | 37,269 | 100.0% (100.0%-100.0%) | 100.0% | 33,302 | 100.0% (100.0%-100.0%) | 97.6% | 37,269 | 100.0% (100.0%-100.0%) | 100.0% | >99.9% |
| cMAF≥1% | Before | 27,295 | 99.9% (99.6%-100.0%) | 98.6% | 27,418 | 100.0% (100.0%-100.0%) | 100.0% | 27,099 | 100.0% (100.0%-100.0%) | 99.9% | 27,419 | 100.0% ( 99.9%-100.0%) | 99.5% | |
| | After | 27,295 | 99.9% (99.6%-100.0%) | 98.6% | 27,418 | 100.0% (100.0%-100.0%) | 100.0% | 27,099 | 100.0% (100.0%-100.0%) | 99.9% | 27,419 | 100.0% (99.9%-100.0%) | 99.5% | 100.00% |
| 200K OQFE-PLINK (n=5,008) | | | | | | | | | | | | | | |

| | | | | | | | | | | | | | | |
|---|---|---|---|---|---|---|---|---|---|---|---|---|---|---|
| All exome variants | Before | 68,005 | 100.0% (98.8%-100.0%) | 92.7% | 80,115 | 100.0% (100.0%-100.0%) | 100.0% | 66,971 | 100.0% (100.0%-100.0%) | 96.2% | 80,115 | 100.0% (100.0%-100.0%) | 99.9% | |
| | After | 65,738 | 100.0% (99.3%-100.0%) | 96.3% | 80,115 | 100.0% (100.0%-100.0%) | 100.0% | 66,969 | 100.0% (100.0%-100.0%) | 96.2% | 80,115 | 100.0% (100.0%-100.0%) | 99.9% | >99.9% |
| cMAF 0%-0.001% | Before | 264 | 0.0% (0.0%-0.0%) | 20.6% | 3,165 | 100.0% (100.0%-100.0%) | 99.6% | 934 | 0.0% (0.0%-0.0%) | 5.4% | 3,165 | 100.0% (100.0%-100.0%) | 100.00 % | |
| | After | 69 | 100.0% (100.0%-100.0%) | 81.2% | 3,165 | 100.0% (100.0%-100.0%) | 99.6% | 934 | 0.0% (0.0%-0.0%) | 5.4% | 3,165 | 100.0% (100.0%-100.0%) | 100.00 % | 98.3% |
| cMAF 0.001%-0.005% | Before | 2,049 | 0.0% (0.0%-100.0%) | 29.8% | 6,989 | 100.0% (100.0%-100.0%) | 99.9% | 1,195 | 100.0% (0.0%-100.0%) | 51.9% | 6,989 | 100.0% (100.0%-100.0%) | 100.00 % | |
| | After | 844 | 100.0% (100.0%-100.0%) | 75.9% | 6,989 | 100.0% (100.0%-100.0%) | 99.9% | 1,195 | 100.0% (0.0%-100.0%) | 51.9% | 6,989 | 100.0% (100.0%-100.0%) | 100.00 % | 99.5% |
| cMAF 0.005%-0.01% | Before | 1,370 | 100.0% (0.0%-100.0%) | 62.6% | 3,050 | 100.0% (100.0%-100.0%) | 100.0% | 1,063 | 100.0% (100.0%-100.0%) | 83.7% | 3,050 | 100.0% (100.0%-100.0%) | 100.00 % | |
| | After | 1,020 | 100.0% (100.0%-100.0%) | 87.5% | 3,050 | 100.0% (100.0%-100.0%) | 100.0% | 1,063 | 100.0% (100.0%-100.0%) | 83.7% | 3,050 | 100.0% (100.0%-100.0%) | 100.00 % | 99.8% |
| cMAF 0.01%-1% | Before | 35,739 | 100.0% (96.9%-100.0%) | 93.0% | 38,328 | 100.0% (100.0%-100.0%) | 100.0% | 35,216 | 100.0% (100.0%-100.0%) | 97.5% | 38,328 | 100.0% (100.0%-100.0%) | 100.00 % | |
| | After | 35,222 | 100.0% (98.0%-100.0%) | 94.9% | 38,328 | 100.0% (100.0%-100.0%) | 100.0% | 35,214 | 100.0% (100.0%-100.0%) | 97.5% | 38,328 | 100.0% (100.0%-100.0%) | 100.00 % | >99.9% |
| cMAF≥1% | Before | 28,583 | 99.9% (99.4%-100.0%) | 99.1% | 28,583 | 100.0% (100.0%-100.0%) | 100.0% | 28,563 | 100.0% (100.0%-100.0%) | 99.9% | 28,583 | 100.0% (99.9%-100.0%) | 99.7% | |
| | After | 28,583 | 99.9% (99.4%-100.0%) | 99.1% | 28,583 | 100.0% (100.0%-100.0%) | 100.0% | 28,563 | 100.0% (100.0%-100.0%) | 99.9% | 28,583 | 100.0% (99.9%-100.0%) | 99.7% | 100.00% |

**Performance of UK BiLEVE Axiom™ array versus whole exome sequencing before and after application of RHA.** Results are split by the WES dataset used as reference -50K FE-VCF and 200K OQFE-PLINK, and by the minor allele frequency calculated from the array before application of RHA (cMAF). We include the following performance metrics: positive predictive value (PPV), negative predictive value (NPV), sensitivity and specificity, as well as the percentage of TP Hets that are retained after applying RHA (only Het genotypes are affected by the algorithm). For each performance metric, the following summary statistics are provided, calculated across all relevant single nucleotide variants: median and inter-quartile range of PPV (the distribution of PPV is not normal) as well as the mean PPV, for the purposes of comparison with Weedon et al. Also included are the number of variants for which the performance metric could be calculated. Variants corresponding to insertions and deletions have been excluded from calculations.

### Supplementary Table D

| Dataset | Before/ After application of RHA | Positive Predictive value (%) | | | Negative Predictive Value (%) | | | Sensitivity (%) | | | Specificity (%) | | | %TP Hets retained after RHA |
|---|---|---|---|---|---|---|---|---|---|---|---|---|---|---|
| | | Number of variants | Median (interquartile range) | Mean | Number of variants | Median (interquartile range) | Mean | Number of variants | Median (interquartile range) | Mean | Number of variants | Median (interquartile range) | Mean | |
| All exome variants | Before | 84,360 | 99.4% (95.9%-99.9%) | 90.5% | 84,801 | 100.0% (100.0%-100.0%) | 100.0% | 83,310 | 100.0% (100.0%-100.0%) | 96.9% | 84,801 | 100.0% (100.0%-100.0%) | 99.9% | |
| | After | 82,867 | 99.7% (98.1%-100.0%) | 95.4% | 84,801 | 100.0% (100.0%-100.0%) | 100.0% | 83,308 | 100.0% (100.0%-100.0%) | 96.9% | 84,801 | 100.0% (100.0%-100.0%) | 99.9% | >99.9% |
| cMAF 0%-0.001% | Before | 2,637 | 33.3% (0.0%-75.0%) | 40.9% | 3,056 | 100.0% (100.0%-100.0%) | 100.0% | 2,217 | 100.0% (0.0%-100.0%) | 70.0% | 3,056 | 100.0% (100.0%-100.0%) | 100.0% | |
| | After | 1,782 | 100.0% (100.0%-100.0%) | 86.3% | 3,056 | 100.0% (100.0%-100.0%) | 100.0% | 2,216 | 100.0% (0.0%-100.0%) | 70.0% | 3,056 | 100.0% (100.0%-100.0%) | 100.0% | 99.9% |
| cMAF 0.001%-0.005% | Before | 8,927 | 66.7% (25.0%-93.3%) | 58.5% | 8,948 | 100.0% (100.0%-100.0%) | 100.0% | 8,329 | 100.0% (100.0%-100.0%) | 87.8% | 8,948 | 100.0% (100.0%-100.0%) | 100.0% | |
| | After | 8,294 | 100.0% (80.0%-100.0%) | 82.8% | 8,948 | 100.0% (100.0%-100.0%) | 100.0% | 8,329 | 100.0% (100.0%-100.0%) | 87.8% | 8,948 | 100.0% (100.0%-100.0%) | 100.0% | 99.9% |
| cMAF 0.005%-0.01% | Before | 3,882 | 91.3% (73.1%-100.0%) | 80.0% | 3,882 | 100.0% (100.0%-100.0%) | 100.0% | 3,865 | 100.0% (100.0%-100.0%) | 94.3% | 3,882 | 100.0% (100.0%-100.0%) | 100.0% | |
| | After | 3,877 | 100.0% (88.9%-100.0%) | 88.3% | 3,882 | 100.0% (100.0%-100.0%) | 100.0% | 3,864 | 100.0% (100.0%-100.0%) | 94.3% | 3,882 | 100.0% (100.0%-100.0%) | 100.0% | 99.8% |
| cMAF 0.01%-1% | Before | 40,017 | 98.9% (96.5%-99.8%) | 95.7% | 40,018 | 100.0% (100.0%-100.0%) | 100.0% | 40,006 | 100.0% (100.0%-100.0%) | 98.4% | 40,018 | 100.0% (100.0%-100.0%) | 100.0% | |
| | After | 40,017 | 99.1% (97.1%-99.9%) | 96.3% | 40,018 | 100.0% (100.0%-100.0%) | 100.0% | 40,006 | 100.0% (100.0%-100.0%) | 98.4% | 40,018 | 100.0% (100.0%-100.0%) | 100.0% | >99.9% |
| cMAF≥1% | Before | 28,897 | 99.9% (99.6%-100.0%) | 99.2% | 28,897 | 100.0% (100.0%-100.0%) | 100.0% | 28,893 | 100.0% (100.0%-100.0%) | 99.9% | 28,897 | 100.0% ( 99.9%-100.0%) | 99.7% | |
| | After | 28,897 | 99.9% (99.6%-100.0%) | 99.2% | 28,897 | 100.0% (100.0%-100.0%) | 100.0% | 28,893 | 100.0% (100.0%-100.0%) | 99.9% | 28,897 | 100.0% ( 99.9%-100.0%) | 99.7% | 100.0% |

**Performance of polymorphic and responsive variants on UK BioBank Axiom™ array versus 200K OQFE-PLINK whole exome sequencing before and after application of RHA.** Results are split by the minor allele frequency calculated from the array before application of RHA (cMAF). We include the following performance metrics: positive predictive value (PPV), negative predictive value (NPV), sensitivity and specificity, as well as the percentage of TP Hets that are retained after applying RHA (only Het genotypes are affected by the algorithm). For each performance metric, the following summary statistics are provided, calculated across all relevant single nucleotide variants: median and inter-quartile range of PPV (the distribution of PPV is not normal) as well as the mean PPV, for the purposes of comparison with Weedon et al. Also included are the number of variants for which the performance metric could be calculated. Variants corresponding to insertions and deletions have been excluded from calculations.

### Supplementary Table E

| Dataset | Before/ After application of RHA | Positive Predictive value (%) | | | Negative Predictive Value (%) | | | Sensitivity (%) | | | Specificity (%) | | | %TP Hets retained after RHA |
|---|---|---|---|---|---|---|---|---|---|---|---|---|---|---|
| | | Number of variants | Median (interquartile range) | Mean | Number of variants | Median (interquartile range) | Mean | Number of variants | Median (interquartile range) | Mean | Number of variants | Median (interquartile range) | Mean | |
| All exome variants | Before | 67,767 | 100.0% (98.8%-100.0%) | 93.0% | 78,470 | 100.0% (100.0%-100.0%) | 100.0% | 65,719 | 100.0% (100.0%-100.0%) | 98.0% | 78,470 | 100.0% (100.0%-100.0%) | 99.9% | |
| | After | 65,639 | 100.0% (99.3%-100.0%) | 96.4% | 78,470 | 100.0% (100.0%-100.0%) | 100.0% | 65,717 | 100.0% (100.0%-100.0%) | 98.0% | 78,470 | 100.0% (100.0%-100.0%) | 99.9% | >99.9% |
| cMAF 0%-0.001% | Before | 195 | 0.0% (0.0%-0.0%) | 22.1% | 2,031 | 100.0% (100.0%-100.0%) | 100.0% | 121 | 0.0% (0.0%-100.0%) | 37.2% | 2,031 | 100.0% (100.0%-100.0%) | 100.0% | |
| | After | 55 | 100.0% (100.0%-100.0%) | 81.8% | 2,031 | 100.0% (100.0%-100.0%) | 100.0% | 121 | 0.0% (0.0%-100.0%) | 37.2% | 2,031 | 100.0% (100.0%-100.0%) | 100.0% | 100.0% |
| cMAF 0.001%-0.005% | Before | 1,913 | 0.0% (0.0%-100.0%) | 30.1% | 6,562 | 100.0% (100.0%-100.0%) | 100.0% | 829 | 100.0% (0.0%-100.0%) | 72.8% | 6,562 | 100.0% (100.0%-100.0%) | 100.0% | |
| | After | 786 | 100.0% (100.0%-100.0%) | 76.8% | 6,562 | 100.0% (100.0%-100.0%) | 100.0% | 829 | 100.0% (0.0%-100.0%) | 72.8% | 6,562 | 100.0% (100.0%-100.0%) | 100.0% | 100.0% |
| cMAF 0.005%-0.01% | Before | 1,341 | 100.0% (0.0%-100.0%) | 63.0% | 2,970 | 100.0% (100.0%-100.0%) | 100.0% | 993 | 100.0% (100.0%-100.0%) | 88.9% | 2,970 | 100.0% (100.0%-100.0%) | 100.0% | |
| | After | 997 | 100.0% (100.0%-100.0%) | 87.9% | 2,970 | 100.0% (100.0%-100.0%) | 100.0% | 993 | 100.0% (100.0%-100.0%) | 88.9% | 2,970 | 100.0% (100.0%-100.0%) | 100.0% | 100.0% |
| cMAF 0.01%-1% | Before | 35,735 | 100.0% (96.9%-100.0%) | 93.0% | 38,324 | 100.0% (100.0%-100.0%) | 100.0% | 35,213 | 100.0% (100.0%-100.0%) | 97.6% | 38,324 | 100.0% (100.0%-100.0%) | 100.0% | |
| | After | 35,218 | 100.0% (98.0%-100.0%) | 94.9% | 38,324 | 100.0% (100.0%-100.0%) | 100.0% | 35,211 | 100.0% (100.0%-100.0%) | 97.5% | 38,324 | 100.0% (100.0%-100.0%) | 100.0% | >99.9% |
| cMAF≥1% | Before | 28,583 | 99.9% (99.4%-100.0%) | 99.1% | 28,583 | 100.0% (100.0%-100.0%) | 100.0% | 28,563 | 100.0% (100.0%-100.0%) | 99.9% | 28,583 | 100.0% ( 99.9%-100.0%) | 99.7% | |
| | After | 28,583 | 99.9% (99.4%-100.0%) | 99.1% | 28,583 | 100.0% (100.0%-100.0%) | 100.0% | 28,563 | 100.0% (100.0%-100.0%) | 99.9% | 28,583 | 100.0% ( 99.9%-100.0%) | 99.7% | 100.0% |

**Performance of polymorphic and responsive variants on UK BiLEVE Axiom™ array versus 200K OQFE-PLINK whole exome sequencing before and after application of RHA.** Results are split by the minor allele frequency calculated from the array before application of RHA (cMAF). We include the following performance metrics: positive predictive value (PPV), negative predictive value (NPV), sensitivity and specificity, as well as the percentage of TP Hets that are retained after applying RHA (only Het genotypes are affected by the algorithm). For each performance metric, the following summary statistics are provided, calculated across all relevant single nucleotide variants: median and inter-quartile range of PPV (the distribution of PPV is not normal) as well as the mean PPV, for the purposes of comparison with Weedon et al. Also included are the number of variants for which the performance metric could be calculated. Variants corresponding to insertions and deletions have been excluded from calculations.

### Supplementary Table F

| Array type | BiLEVE | BiLEVE | UK Biobank |
|---|---|---|---|
| Cohort | 50K WES | 200K WES | 200K WES |
| Which BRCA variants? | Weedon [1] n = 1076 | Weedon[1] n = 99 | cMAF < 0.01% n = 460 |
| TPs | 5 | 6 | 2529 |
| FNs | 0 | 1 | 454 |
| Exome Het Array NoCall | 2 | 4 | 809 |
| Overall Sensitivity | 100% | 85.7% | 84.8% |
| Variants with 100% sensitivity | 5 of 5 | 5 of 6 | 318 of 388 |
| Pre-RHA FPs | 742 | 132 | 2211 |
| Overall PPV | 0.67% | 4.4% | 53.4% |
| Post-RHA FPs | 344 | 67 | 598 |
| Overall PPV | 1.4% | 8.2% | 80.9% |
| TPs removed by RHA | 0 | 0 | 3 (0.12%) |
| FPs removed by RHA | 53.6% | 49.2% | 73.0% |

**Characterization of BRCA variant performance on the Axiom™ arrays used by UK Biobank.** The variants studied by Weedon et al. [1] were graciously provided by the authors, and the data was re-analyzed to include only those variants that were present on the specified array. Each data point represents one Het call in one sample. Some participants had more than one Het call.

To provide additional data from the UK Biobank array, we also present results from all rare variants (cMAF < 0.01%) within the transcribed regions of *BRCA1* and *BRCA2*.
